# Automating Assessment of the Undiscovered Biosynthetic Potential of Actinobacteria

**DOI:** 10.1101/036087

**Authors:** Bogdan Tokovenko, Yuriy Rebets, Andriy Luzhetskyy

**Affiliations:** Actinomycetes Metabolic Engineering Group, Helmholtz Institute for Pharmaceutical Research Saarland, Saarbrücken, Germany; Department of Pharmaceutical Biotechnology, Faculty of Natural Sciences and Technology, University of Saarland, Saarbrücken, Germany

**Author notes:** Corresponding author (AL), (BT).

## Abstract

**Background**. Biosynthetic potential of Actinobacteria has long been the subject of theoretical estimates. Such an estimate is indeed important as a test of further exploitability of a taxon or group of taxa for new therapeutics. As neither a set of available genomes nor a set of bacterial cultivation methods are static, it makes sense to simplify as much as possible and to improve reproducibility of biosynthetic gene clusters similarity, diversity, and abundance estimations.

**Results**. We have developed a command-line computational pipeline (available at https://bitbucket.org/qmentis/clusterscluster/) that assists in performing empirical (genome-based) assessment of microbial secondary metabolite gene clusters similarity and abundance, and applied it to a set of 208 complete and de-duplicated Actinobacteria genomes. After a brief overview of Actinobacteria biosynthetic potential as compared to other bacterial taxa, we use similarity thresholds derived from 4 pairs of known similar gene clusters to identify up to 40-48% of 3247 gene clusters in our set of genomes as unique. There is no saturation of the cumulative unique gene clusters curve within the examined dataset, and Heap’s alpha is 0.129, suggesting an open pan-clustome. We identify and highlight pitfalls and possible improvements of genome-based gene cluster similarity measurements.

## Introduction

Theoretical estimates of Actinobacteria biosynthetic potential often point at a huge possible number of unique antibiotics which are still to be discovered [1-3]. Such an estimate is indeed important as a test of further exploitability of a taxon for new therapeutics. Focus on the *Streptomyces* genus is quite understandable: of the circa 12000 secondary metabolites with antibiotic activity known in 1995, 55% were produced by *Streptomycetes* and additional 11% by other Actinobacteria [4].

While Actinobacteria were *the* sources of new antibiotics in the 1960-70s, progressively fewer and fewer new antibiotics were found since then, feeding a growing disappointment and disbelief in the ability to find more novel compounds. However, it is now recognized [5,6] that most of the BGCs (biosynthetic gene clusters) are silent and require activation to produce compounds. Thus, genome-based estimate, which also includes silent BGCs, is important for the revival of interest and belief in the potential of Actinobacteria.

Currently dominating antibiotics discovery strategy is to search wide – that is, characterize and sequence more strains (for example, marine Actinobacteria), looking for promising compounds. Quite apparently such a strategy is not cheap, and normally it does not overcome the issue of silent BGCs, as these remain silent unless specific steps are taken to activate them. BGC activation is a laborious process. Therefore, an important step preceding BGC activation experiments is prioritisation of the most promising genomes and clusters.

Moreover, based on our own experience of understanding and working with BGCs, we felt that there is a need to quantify a persistent feeling of finding highly similar BGCs in different genomes. Thus, we decided to perform medium-scale genomes-based estimation of the number of unique BGCs discovered with each new genome, as well as develop and make public the tools for comparing BGCs en masse.

With this study we prepare an analysis-driven strategy for estimating BGCs diversity/similarity in a sample of multiple genomes, and also (to a lesser degree) a strategy for BGCs (and genomes) prioritisation before their activation. Ideally, this strategy would start with an automated exploration of a large corpus of BGCs (looking for similarities and/or specific genes), then proceed through manual examination (and, possibly, adjusting annotations) of the candidate BGC(s), and culminate in the experimental activation of the candidate BGC(s). Based on our genome set, we answer a number of questions about the secondary metabolites genomics of Actinobacteria. Do highly similar BGCs occur in different genomes? How often? How many previously unseen, novel BGCs one may expect when sequencing new strain’s genome? Which BGC types are the least and the most unique? What should we expect in terms of novel BGCs if we extrapolate from the current set of genomes?

There are several significant benefits of automated estimation compared to infrequent individual studies. First of all, estimation results become more easily reproducible. Secondly, comparison results become readily available as a simple table, which only requires re-calculations when new genomes/clusters are added. Finally, it is much easier (and faster) to update estimates and comparisons as new genomes become available, including discoveries of new species, or development of new bacteria cultivation methods, significantly extending the range of microorganisms which can be grown and studied in the lab.

To the best of our knowledge, this is the first attempt at automating empirical (genome-based) quantification of Actinobacteria potential. Presented approach and software can likely be applied to other natural sources of BGCs.

## Results and discussion

### Overview of potential

Why do we focus on Actinobacteria? Empirically, Actinobacteria were the largest natural product source since 1960s. But does this taxon also stand out at the genomic level, taking into account silent clusters? To answer this question, we performed an overview study of 2775 bacterial genomes. We had also performed the same overview separately for 285 genomes of Actinobacteria.

We wanted to know how many putative BGCs (biosynthetic gene clusters) can be identified using antismash2 in the genomes, and then group this data by genus/family and other taxonomic levels to reveal highest-cluster-count groups.

Prior to the overview analysis highly similar genomes and genomes shorter than 2.5 megabases were removed as described in the *Methods*. Among the smallest genomes, the following phyla were highly represented (roughly in the order of increasing genome size): Proteobacteria, Tenericutes, Bacteroidetes, Spirochaetes, Chlamydiae, Chloroflexi, Firmicutes, Euryarchaeota, Crenarchaeota, Korarchaeota, Thaumarchaeota, Aquificae, Cyanobacteria, Thermotogae. There were also a few Actinobacteria (e.g. *Tropheryma, Cryptobacterium, Bifidobacteria*, etc). The highest number of BGCs among genomes smaller than 2 megabases was observed in Cyanobacteria - up to 13 gene clusters in *Prochlorococcus marinus str*. MIT 9313. Table S1 contains the spreadsheet summary of small genome characteristics.

The two largest genomes were *Sorangium cellulosum* So0157-2 (NC_021658) and *Sorangium cellulosum* So ce56 (NC_010162) at 14.8 and 13 megabases long, respectively. The next-largest genome was *Streptomyces rapamycinicus* NRRL 5491 (NC_022785) at 12.7 megabases.

Among Actinobacteria, the Actinomycetales order (Actinobacteriadae subclass) is the most well-represented (232 genomes), followed by Bifidobacteriales (33 genomes, the entire Bifidobacteriaceae suborder of this dataset). The most well-represented suborder is Corynebacterineae (127 genomes), followed by Bifidobacteriaceae (33 genomes), Micrococcineae (30 genomes), Streptomycineae (19 genomes) and others. *Mycobacterium* (63 genomes), *Corynebacterium* (50 genomes), and *Bifidobacterium* (30 genomes) dominate at the genus taxonomical level.

Actinobacteria had an average of 10.7 BGCs per genome (median 6), while the complete set of genomes had mean 5.6 (median 4). Maximal number of BGCs identified was 50 in *Streptomyces bingchenggensis* BCW-1 (NC_016582). Fig. 1 (top) shows boxplots of BGC counts per genome for the full genome set and Actinobacteria genomes.

**Fig. 1.**
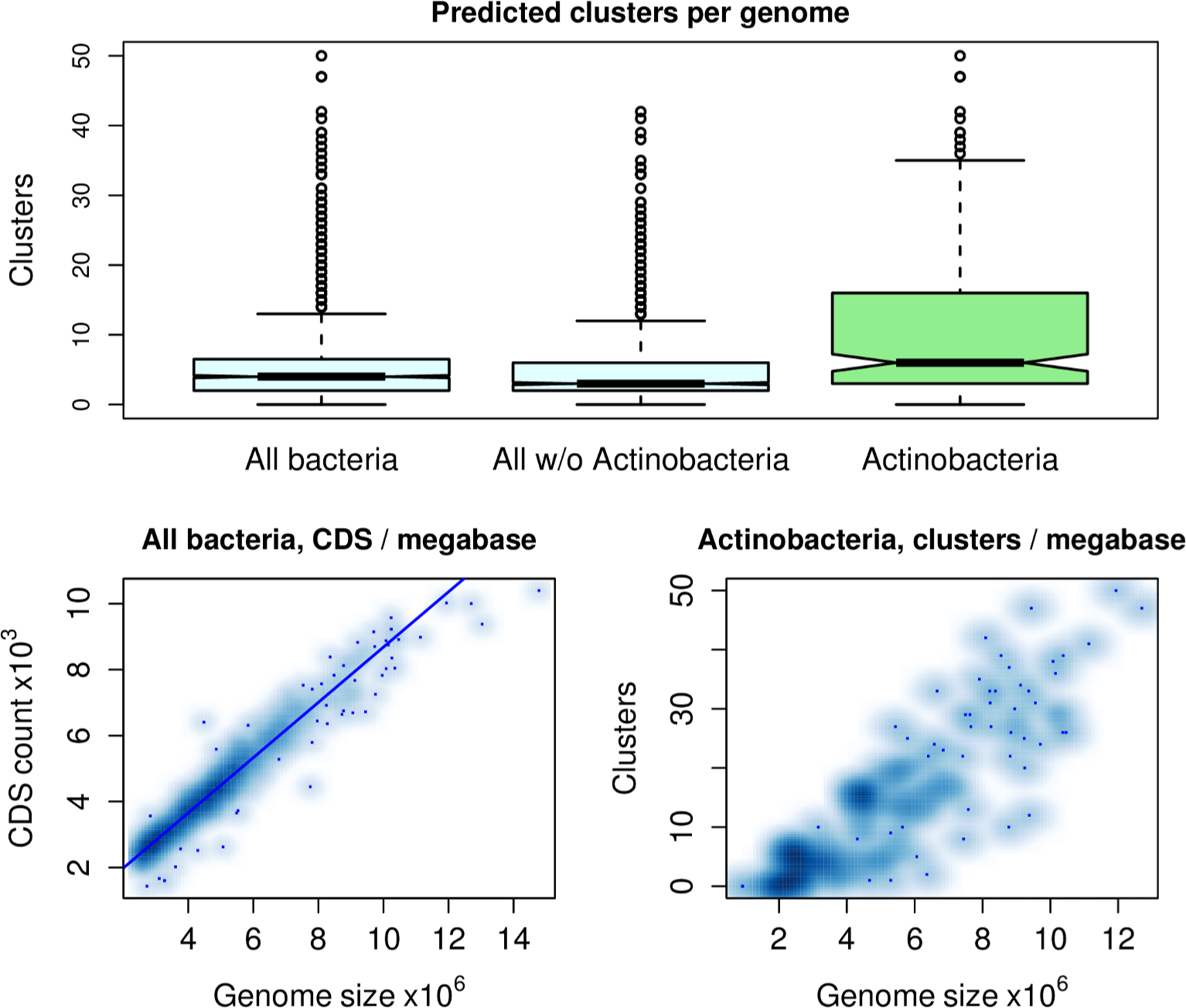
Predicted gene clusters per genome. **Top:** boxplots of predicted gene cluster counts per genome for 1651 bacterial genomes larger than 2.5 megabases, 1508 non-Actinobacteria large genomes, and for 278 Actinobacteria genomes of all sizes. **Bottom left:** smoothed scatterplot of CDS counts and genome sizes for 1651 bacterial genomes. **Bottom right:** smoothed scatterplot of putative gene cluster counts and genome sizes for 278 Actinobacteria genomes.

There is a clear linear relationship between genome size and the number of annotated CDS (Fig. 1, bottom left; Pearson correlation 0.97, p-value < 10^−5^). Expressed as a linear model, `CDScount = 304.3 + 837.8xgenome_size_in_megabases`. Linear relationship holds for all studied subsets of bacterial genomes.

In Actinobacteria the count of putative BGCs appears to correlate better with genome size (Fig. 1, bottom right; Pearson correlation coefficient 0.88, p-value < 10^−5^) than in all bacteria (not shown). Expressed as a linear model, `clusters_count = −6.8 + 3.9 × actino_genome_size_in_megabases` (almost 4 clusters per megabase, except for the first 1.74 megabases).

At the phylum taxonomy level, the two leaders are Actinobacteria and Proteobacteria. Cyanobacteria are also promising: this phylum has at least 3 genomes with over 20 putative BGCs each (Fig. 2). Fig. 2 shows that the majority of Proteobacteria have fewer than 10 BGCs (mean is 4.9 clusters/genome), but there is also a group of high BGC count outliers which make this phylum stand out. Cyanobacteria mean is higher than that of Proteobacteria (8.5 clusters/genome). Going down to the class level, we see that specifically Beta- and to a lesser extent Delta-Proteobacteria have the most promising BGCs per genome count (not shown).

**Fig. 2.**
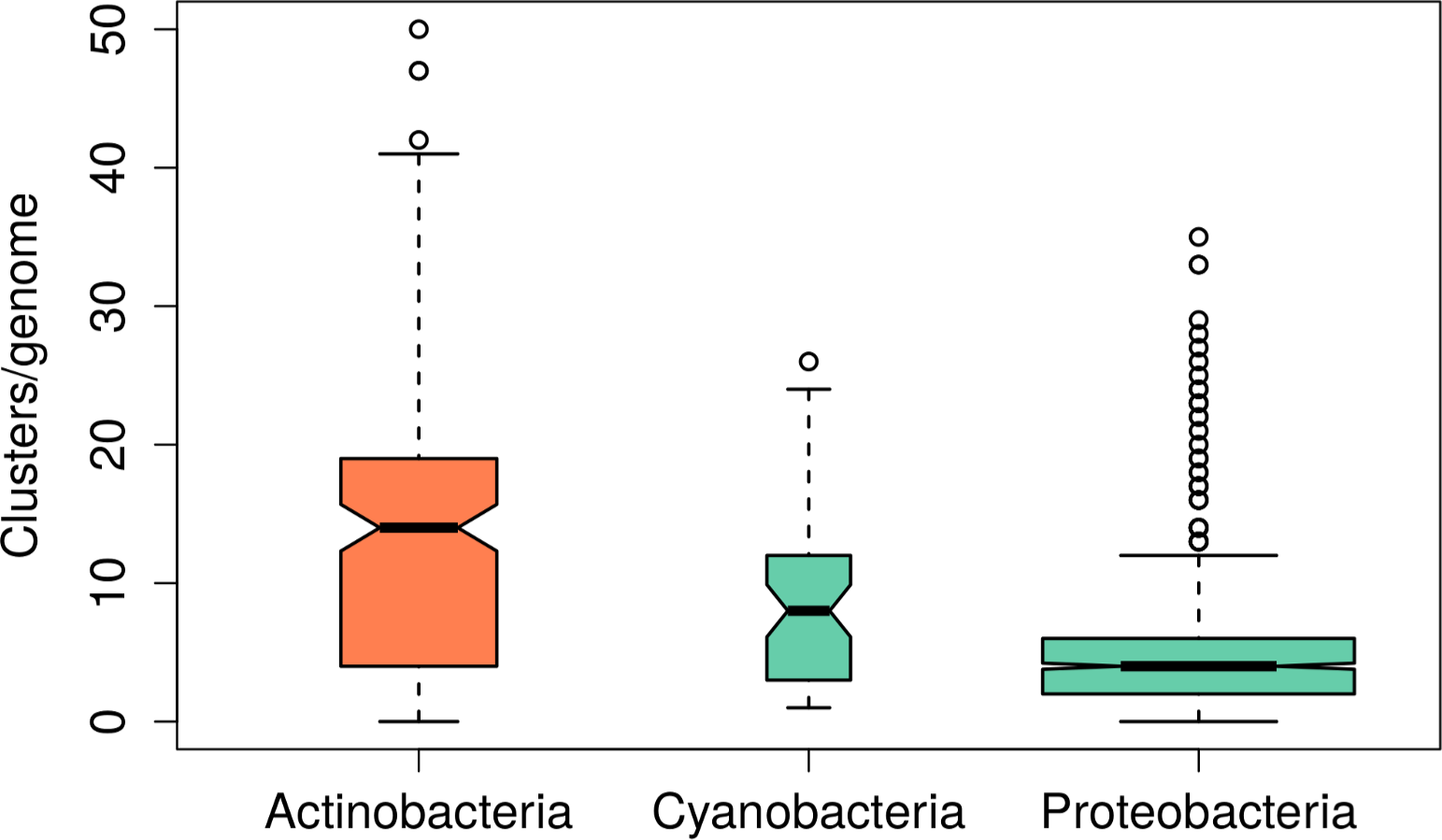
Boxplot of putative gene clusters count per genome by phyla. Only phyla with at least 9 genomes, and with mean gene clusters per genome larger than or equal to 4 were included. Width of each boxplot is proportional to the square root of the number of genomes in the phylum.

At the genus level (Fig. 3), *Streptomyces* clearly dominate. Runners-up are *Frankia* and *Rhodococcus* (both with low genome counts), also *Burkholderia* and *Mycobacteria*. 60 *Mycobacteria* genomes appear to be fairly heterogeneous with respect to their BGCs count per genome, as can be seen by the highest number of outliers this genus has in Fig. 3.

**Fig. 3.**
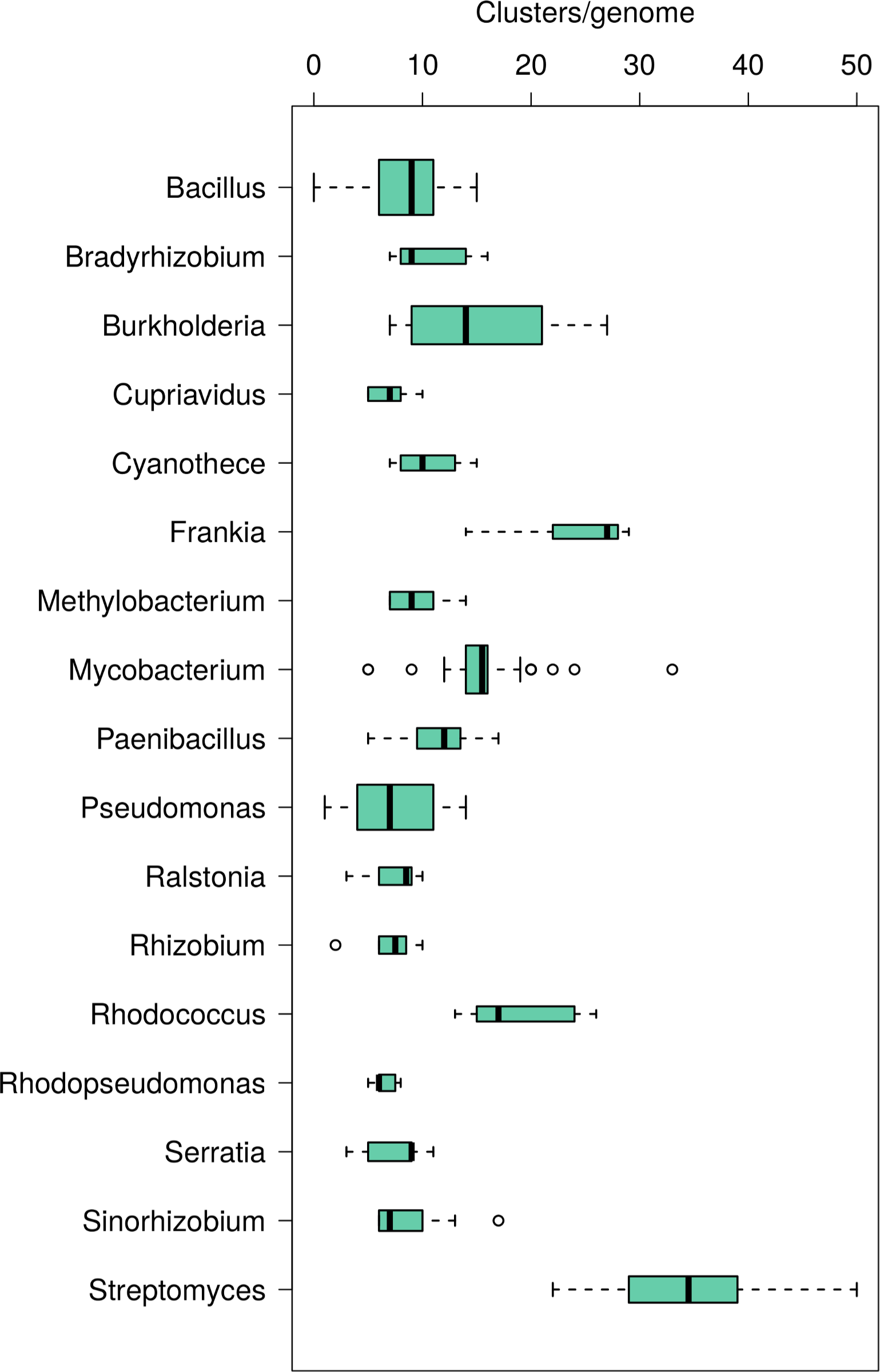
Boxplot of putative gene clusters count per genome by genus. Only genii with at least 5 genomes, and with mean gene clusters per genome larger than or equal to 6 were included. Width of each boxplot is proportional to the square root of the number of genomes in the genus.

For a more in-depth analysis of BGCs by their types and proportions within Actinobacteria please see [7].

### Automating biosynthetic potential assessment

After confirming that Actinobacteria indeed have the highest genomic potential to produce bioactive compounds, the next step is to assess BGCs similarity. We developed our method based on how a human expert would approach the task of measuring BGCs similarity. A pair of (potentially similar) BGCs is the basic analysis unit of our method. A normalized BGC similarity score enables multiple types of downstream analysis. An ability to calculate this similarity score from different metrics ensures some flexibility of the developed pipeline to handle multiple and changing scenarios.

The software pipeline comes in two parts: the primary genomes processing Python script (with multiple external dependencies), which produces a CSV file for the second part – R scripts for CSV data analysis and summarization (Fig. S10 and dataset S11). The software has command-line interface and was tested on Linux only. However, program options and the source code are well-documented. Source code is available at <u>https://bitbucket.org/qmentis/clusterscluster</u>.

Python program accepts a list of (possibly multi-locus) GenBank (.gb or .gbk) files, 1 per genome. As long as these GenBank files can be read with BioPython [8], they will be compatible with the program. GenBank was chosen as a sufficiently common sequence and annotation format, though annotations are currently not required for processing. In fact, only genomic sequence(s) and sequence meta-data (ID, species/strain information) are used. Exact and ID-only duplicates are detected and reported by the program.

The BGC prediction software that we used, AntiSMASH 2 [9], can either use existing gene annotations or quickly predict CDS using Glimmer. As gene annotations can be quite different, affecting BGC prediction, we decided to strip any existing gene annotations and allow AntiSMASH 2 to use Glimmer for all genomes. This likely had a negative effect on the quality of BGC prediction, but also removed any annotation quality biases.

AntiSMASH 2 works by identifying core secondary metabolite genes, and then adds variable-size extensions (5-20kbp, depending on BGC type) up- and down-stream from the core genes. These extensions contain both cluster genes and some non-cluster genes, possibly hampering cluster similarity analysis. We provide an optional patch which allows using AntiSMASH 2 without these extensions. However, as tailoring enzymes are important, by default BGC extensions are included. All the CDS sequences are then translated to a multi-fasta protein file, which is then used for all protein-level comparisons.

Our software can identify cluster genes from different genomes which belong to the same group of orthologs across all the genomes in the analysis. We use InParanoid [10] and QuickParanoid (<u>http://pl.postech.ac.kr/QuickParanoid/</u>) for orthologous groups construction across all the analyzed genomes. InParanoid was slightly modified to allow incremental addition of genomes without recalculating existing genome pair similarities. This processing step generates a single table of all orthologous groups for the analyzed genomes. Genes from different clusters belonging to a common orthologous group form a single “link” between the BGCs; the number of such orthologous links can then be used to quantify cluster similarity.

In our earlier pilot study of BGCs similarity (not published) we found that gene orthology is not powerful enough for BGC comparisons. For example, many NRPS proteins despite performing different synthesis steps have high (>70%) identities, which assigns them all to a handful of orthologous groups. Orthology analysis, which requires 4 full-genome blasts per one pair of genomes, is also quite computationally intensive. To partially remedy this, we provide a separate script dependent only on a local InParanoid directory to allow splitting computations to multiple computers/nodes/GRID workers. Although we keep the orthologous groups functionality, our focus is now on full-length protein-protein alignments, which provide better resolution (lower granularity) of gene similarity measurements.

Full length protein alignments are performed in parallel with usearch [11], an extremely fast heuristic protein alignment/search tool. It is possible to avoid heuristics by enabling the full dynamic programming mode of usearch, which is significantly slower. In this initial version, all possible pairs of proteins from BGCs are aligned. Computing time for this step can be significantly improved in future versions by only aligning reasonable pairs (e.g. those with similar lengths). All protein alignment results are saved to a file-based cache, enabling process restarts and incremental addition of genomes to the analysis.

For all genes similar between any two clusters, their average protein identity is calculated, and gene order in both clusters is compared. Finally, all collected and computed results are written to a single CSV file (one similar BGC pair per row, dataset S11) for further analysis. See Fig. S10 for pseudo-code of BGCs comparison, as a part of genomes processing flowchart.

Further analysis and figure plotting are performed with several R scripts, which were used for the preparation of this manuscript.

### Gene cluster similarity definition

Ideally, two BGCs should be referred to as similar if the compounds they produce are similar. If one had compound structures for both BGCs, one could then use Tanimoto compound similarity index with a certain threshold for this purpose. However, it is not immediately obvious what should the value of such a threshold be: indeed, what is the minimal difference between similar and dissimilar molecules? For example, would methylation make a new molecule dissimilar? What about one more methylation? Moreover, predicting compound structure purely from gene composition of a cluster is neither precise nor reliable. This further complicates the definition of what “similar gene clusters” really are (except for obvious cases, like our control BGCs). Although we thus realize that both the metric and the threshold we are using are not perfect, we believe they provide useful proxies for identifying similar and unique BGCs, responsible for biosynthesis of small molecules with new chemotypes. Making our toolbox available allows interested parties to modify and create new metrics and thresholds, adapting them to specific tasks.

### Gene cluster similarity data table

After developing the software and being motivated to test it, we sought applying it and getting BGCs similarity estimate for a subset of Actinobacteria genomes. In this and the following sections we describe some of the analyses of the data generated with our software.

Using 208 thoroughly de-duplicated complete Actinobacteria genomes as inputs, we obtained a data table of BGC pairs and their similarity. This table is the starting point for all the following analyses. The resulting spreadsheet is available as dataset S11. Columns have the following meanings:

- **is_intra:** “1” means that both BGCs of the pair belong to the same genome (intra-genome similarity); “0” means inter-genome similarity;
- **genome1_ID, genome2_ID:** GenBank accession number for the longest nucleotide sequence of the genome (i.e. chromosome); for a subset of non-GenBank genomes – an internally assigned genome ID;
- **speciesl, species2:** species information extracted from the original GenBank files; this is missing for some of the non-GenBank genomes;
- **clusterl, cluster2:** numeric identifiers of BGCs within the genome, as assigned by the cluster prediction method;
- **typel, type2:** generic BGC type, as assigned by the cluster prediction method;
- **genes1_count, genes2_count:** number of genes in the 1^st^ and 2^nd^ clusters of the pair;
- **size1_kb, size2_kb:** BGC sizes in kilobases;
- **ortholinks_count1, ortholinks_count2:** number of orthology links between the BGCs (see *Methods* for details); we have not calculated these for our test set;
- **similar_genes_count:** a number of unique gene pairs between the clusters with protein identity above 60%;
- **avg_protein_identity:** actual average protein identity of all similar genes;
- **P, K, S:** Pearson, Kendall, and Spearman correlations of two integer vectors of similar gene positions in both clusters;
- **ratio1, ratio2:** ratio of similar_genes_count to genes1_count and genes2_count, respectively;
- **ratioa:** an average of ratio1 and ratio2;
- **clusterid1, clusterid2:** string concatenation of genome1_ID with clusterl and genome2_ID with cluster2, respectively; this serves as a unique BGC identifier.

Of these columns, the last five (ratio1, ratio2, ratioa, clusterid1, clusterid2) were added by the additional preprocess.R script.

### Gene-level synteny

To quantify local gene-level synteny for BGC pairs, we tested as measures Spearman and Kendall rank-correlation coefficients, as well as Pearson’s correlation coefficient and Kendall tau distance for two vectors of similar gene positions within their respective gene clusters. None of these methods was perfect for gene synteny. Rank-correlation measures ignore gene deletions/insertions, while Pearson’s correlation is not sensitive to gene rearrangements in longer gene clusters. We are working on a more sensitive gene-level synteny metric, which will be added to the software repository after testing.

### Thresholds and unique gene clusters

As already mentioned in *Gene cluster similarity definition*, defining a single perfect threshold might not be possible, at least for genome-based gene cluster similarity comparison.

Similarity thresholds that we used are primarily based on values obtained for 4 pairs of known BGCs from different genomes responsible for the production of the same or highly similar compound: landomycin, microsclerodermin, moenomycin, nonactin (Table 1). First, only BGC sequences were compared, out of their genomic contexts - that is, with well-defined boundaries (manually annotated, no extensions with unrelated genes; top 4 rows of Table 1). For moenomycin, complete genome sequences were also available, so the manually annotated BGCs were first compared to their respective automatically-annotated counterparts in the genomic context (yielding lower similarity scores), and then both S. *Ghanaensis* and S. *Clavuligerus* complete genomes were analysed, yielding the lowest control similarity score of 0.29 for automatically annotated moenomycin BGCs in their genomic context (last 3 rows of Table 1). Based on these controls, the lowest acceptable BGC similarity score was set to 0.5 - sufficient for identifying all but one of control cases as similar.

**Table 1.** Pairs of previously characterized similar gene clusters.

| Gene cluster | Species | Gene pairs | Similarity score |
| --- | --- | --- | --- |
| Landomycin | *S. cyanogenus* / *S. Globisporus* | 26 | 0.79 |
| Microsclerodermin | 2 *Streptosporangium* species | 9 | 0.53 |
| Moenomycin | *S. ghanaensis* / *S. Clavuligerus* | 12 | 0.73 |
| Nonactin | *S. sp. CcalMP-8W* / *S. Fulvissimus* | 24 | 0.84 |
| Moenomycin | *S. clavuligerus* cluster/genome | 14 | 0.55 |
| Moenomycin | *S. ghanaensis* cluster/genome | 15 | 0.67 |
| Moenomycin | *S. ghanaensis* / *S. Clavuligerus* whole genomes | 13 | 0.29 |

Looking into the 0.29 similarity score for moenomycin in genomic context, we found that a nearby NRPS-T1PKS BGC and a single T1PKS gene in *S. Clavuligerus* were appended to the predicted moenomycin BGC, resulting in a single huge composite cluster (97 kbp and 85 genes versus 35 kbp and 30 genes for predicted moenomycin in *S. Ghanaensis*). Thus, even despite 13 similar genes with average protein identity 68.3%, these two gene clusters received a low similarity score. In our method, the estimated percentage of unique BGCs will be sensitive (proportional) to the frequency of predicting a single combined/composite BGC instead of two or more distinct clusters.

Despite using control BGCs, the 0.5 similarity score would benefit from more supporting evidence. Thus, we deemed it necessary to estimate the variability of unique BGCs percentage over the full range of threshold values (Fig. 4). Between similarity score threshold values 0.2 and 1.0 the percentage of unique BGCs grows slower than the threshold (slope < 1), indicating relative stability of the unique percentage estimate, and suggesting that minimal percentage of unique BGCs in our dataset is approximately 40 (at threshold value 0.2).

**Fig. 4.**
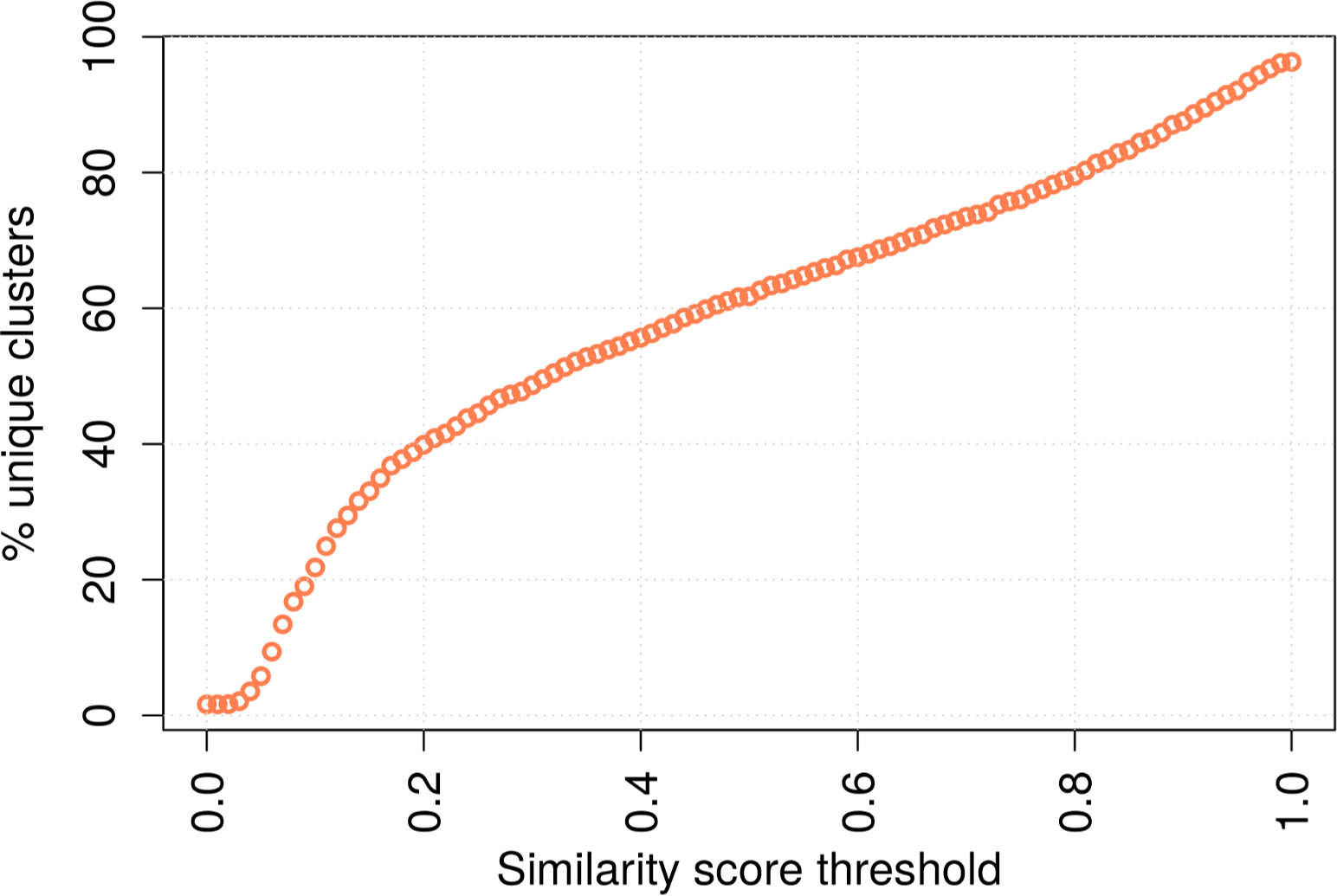
Effects of changing similarity score threshold on the percentage of unique gene clusters.

To summarize, we provide the following support for the 0.5 score threshold used throughout this manuscript:

- it is the lowest threshold allowing detection of 4 pairs of control BGCs (outside their genomic context and after manually refining BGC boundaries);
- it is higher than the minimal threshold 0.2 suggested by diagnostic plot for thresholds (Fig. 4).

The 0.5 threshold is too high to detect the only control BGC tested in its genomic context (without manual cluster boundaries adjustment). This suggests that our estimates of unique BGCs may have upward bias.

### Gene clusters diversity: percentage of unique clusters

After building the BGC similarity table, we focused on finding the percentage of unique BGCs – that is, clusters which do not have above the threshold similarities to other clusters. We decided to use the *ratioa* column as the primary measure of BGC similarity (hereafter referred to as similarity score). It is of course possible to use other dataset columns and their combinations as measures of similarity; however, existing analysis R scripts will need to be updated for that.

Based on our control BGCs (see *Thresholds and unique gene clusters* above), we started with cluster similarity score 0.5 or higher (threshold 1), identifying 61.7% BGCs as unique. This threshold seems low, but 4 pairs of control similar BGCs would need such a low score to be identified as similar pairs. Low score may identify some of the longer unique BGCs as “similar”, but given the presence of up- and down-stream extensions around putative BGCs, it is more likely that even such a low threshold would fail to detect truly similar BGCs, as exemplified by the low similarity score of 0.29 for the control moenomycin BGC in 2 different genomic contexts (Table 1 in *Thresholds and unique gene clusters*).

If, in addition to threshold 1, we require at least 10 genes to be similar (protein identity > 60%) between the BGCs (threshold 2), then the percentage of unique clusters increases to 73%. The number of genes for threshold 2 was also based on our control clusters (Table 1) and diagnostic plots for thresholds (Appendix S2). This additional threshold has two major effects: first, clusters shorter than 10 genes are automatically considered “unique”; second, shorter spurious similar BGCs obtained at similarity score threshold 0.5 are filtered out. Based on this threshold increase, we can conclude that clusters shorter than 10 genes constitute less than 11 % of all the BGCs, and that false-positive detections of shorter similar BGCs with a score threshold 0.5 are smaller than 11%. Examining distribution of putative BGCs by predicted type in 3 sets – all, unique, and similar – shows that additional 10-gene threshold affects the most bacteriocin, butyrolactone, ectoine, melanin, and siderophore BGCs, which are indeed normally shorter than 10 genes (Fig. 5). Taken together, decrease of the count of these BGC types explains almost all of the estimate change. In other words, requiring at least 10 genes to be similar does not change the unique BGCs estimate based on similarity score.

**Fig. 5.**
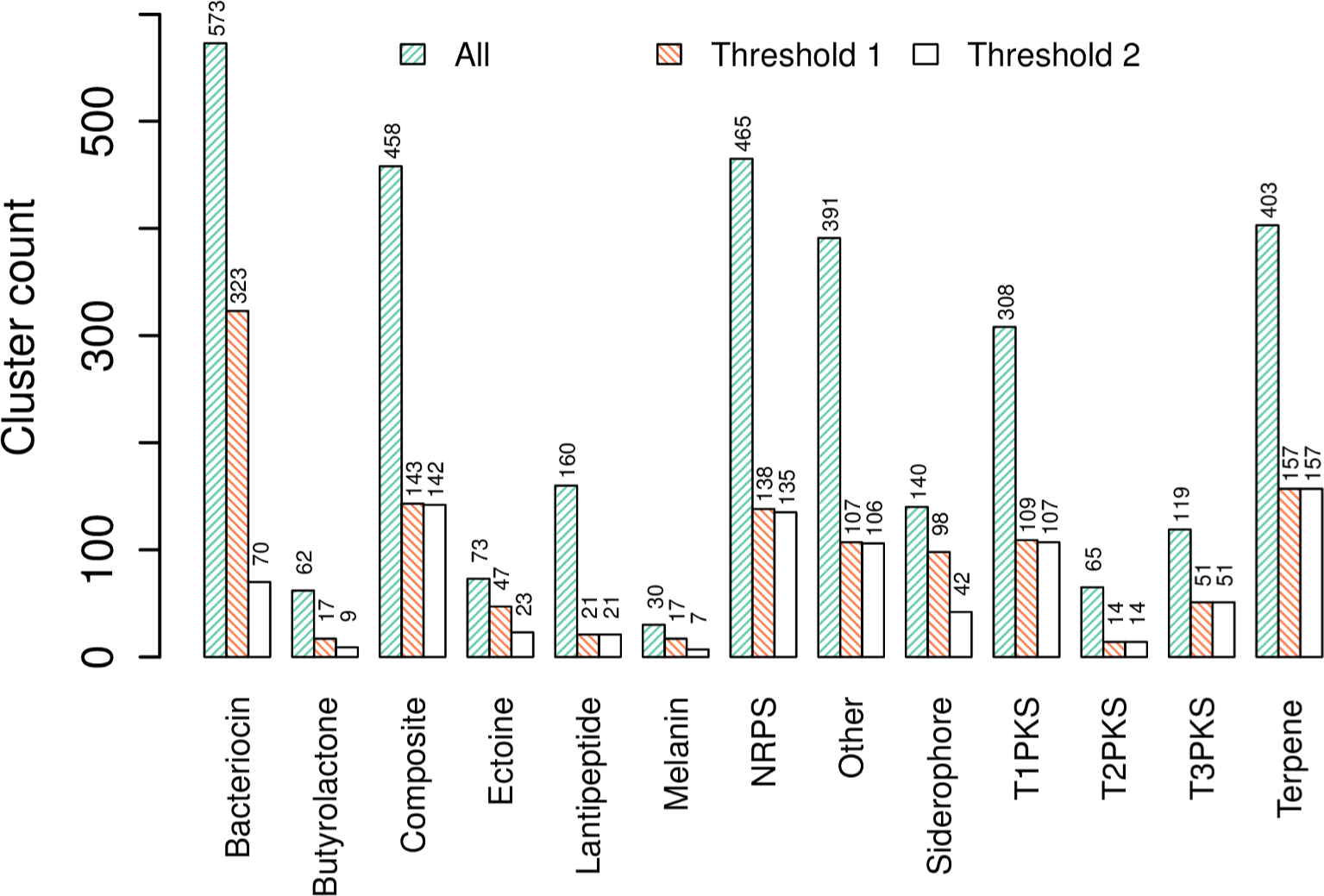
Distribution of secondary metabolite gene clusters by type in 3 datasets: all 3247 gene clusters, non-unique at threshold 1 (similarity score above 0.5), non-unique at threshold 2 (similarity score above 0.5 and at least 10 similar genes).

Interestingly, if we only use the existence of 10 similar genes as a threshold (disregarding all the other metrics of BGC pair similarity), we get 63.35% of unique BGCs (see Appendix S2 for a diagnostic plot of similar genes threshold versus percent unique BGCs). Adjusting by 11% of the clusters known to be shorter than 10 genes, we arrive at the base estimate of 52% unique BGCs.

As we learn from diagnostic plots, around 4% of all BGCs with near-perfect similarities might come from the undetected similar genomes, decreasing our base estimate from 52% to 48%.

To summarize, we think that the 40-48% range of unique BGCs is a good estimate of uniqueness for the 3247 BGCs examined in 208 de-duplicated genomes.

### Highest counts of unique gene clusters per genome

The next question we wanted to answer is which genomes have the highest unique BGC counts (as defined in the previous section). After sorting all the genomes by the descending count of unique BGCs, we obtain a list of soil-dwelling Streptomycetacea (Table 2). The only surprising entry here is *Nocardia brasiliensis*, which can be pathogenic. As we know, the larger the automatically predicted BGC is – possibly including several unrelated smaller clusters – the lower is the chance that it will be properly recognized as similar to something else. Thus, this table of the most unique BGCs-rich genomes is likely to be biased towards bigger or closely located but unrelated BGCs.

**Table 2.** Top 5 genomes by absolute counts of unique secondary metabolite gene clusters.

| Genome ID | Name | Family | Habitat | Unique gene clusters | Total gene clusters |
| --- | --- | --- | --- | --- | --- |
| CP007155.1 | *Kutzneria albida*<br>DSM 43870 | Pseudonocardiaceae | Soil | 53 | 54 |
| NC_016582.1 | *Streptomyces bingchenggensis*<br>BCW 1 | Streptomycetaceae | Soil | 46 | 53 |
| NC_016109.1 | *Kitasatospora setae* KM-6054 | Streptomycetaceae | Soil | 43 | 49 |
| NC_018681.1 | *Nocardia brasiliensis*<br>ATCC 700358 | Nocardiaceae | Soil | 40 | 51 |
| CM001015.1 | *Streptomyces clavuligerus*<br>ATCC 27064 | Streptomycetaceae | Soil | 40 | 52 |

We had previously reported that secondary metabolism genes occupy a significantly bigger proportion of *K. albida* genome compared to other actinomycetes, with 9.6% of all genes located inside manually-curated secondary metabolite BGCs [12]. It is thus not too surprising to find *K. albida* genome at the top of Table 2. But does it really have 53 unique BGCs? Expert examination of 46 BGCs reported previously [12] identified at least 9 BGCs which are not unique (a whiE-like type II PKS, several siderophores and terpenes, and also bacteriocin and ectoine BGCs). It is thus clear that 53 unique BGCs is an overestimate. Correspondingly, the above-derived 40-48% of unique BGCs estimate should be interpreted as an upper bound estimate.

### Gene clusters diversity: groups of gene clusters

Another way to look at gene clusters diversity is to group/cluster them by similarity. We provide an R script which does this based on the similarity score as defined above (see *Methods/Grouping gene clusters by similarity*). Applying this approach at similarity score threshold 0.5, we identified 2393 groups of 3247 BGCs.

Doroghazi *et al* [2] used a clustering-based approach to identify groups of similar BGCs. In our study a single group of clusters contains on average 1.4 clusters, while groups identified by Doroghazi and colleagues have 2.8 clusters per group (average calculation takes into account singleton groups). Apparently, the similarity threshold that we use is more stringent than in the aforementioned study. In addition, Doroghazi *et al* used their own method for finding clusters; despite having much higher fraction of clusters-rich Streptomyces (approximately 40% of all genomes examined, compared to our 15%), Doroghazi *et al* found on average 13.8 clusters per genome, compared to 15.9 found with AntiSMASH 2 in this study.

### Extrapolating the number of unique gene clusters

Estimation of gene cluster similarity would be incomplete without extrapolating to more genomes. A popular approach is to apply rarefaction analysis. However, rarefaction analysis was originally designed for interpolating to a smaller sample size, not extrapolating to a larger sample size (at least without assuming a specific underlying distribution), and it is not clear if all the assumptions are met (such as sufficiently large sample size and random distribution of “individuals” in samples), in addition to the underlying biological limitations (see *Rarefaction analysis* below).

Thus, we had applied a more appropriate resampling approach to look for saturation of the total unique BGCs curve (Fig. 6). With a little over 200 genomes, the curve is slightly bent, but there is definitely no saturation. We thus refrained from extrapolating this curve to higher numbers of genomes. See Appendix S3 for 2 more scatterplots, at thresholds 0.2 and 0.8.

**Fig. 6.**
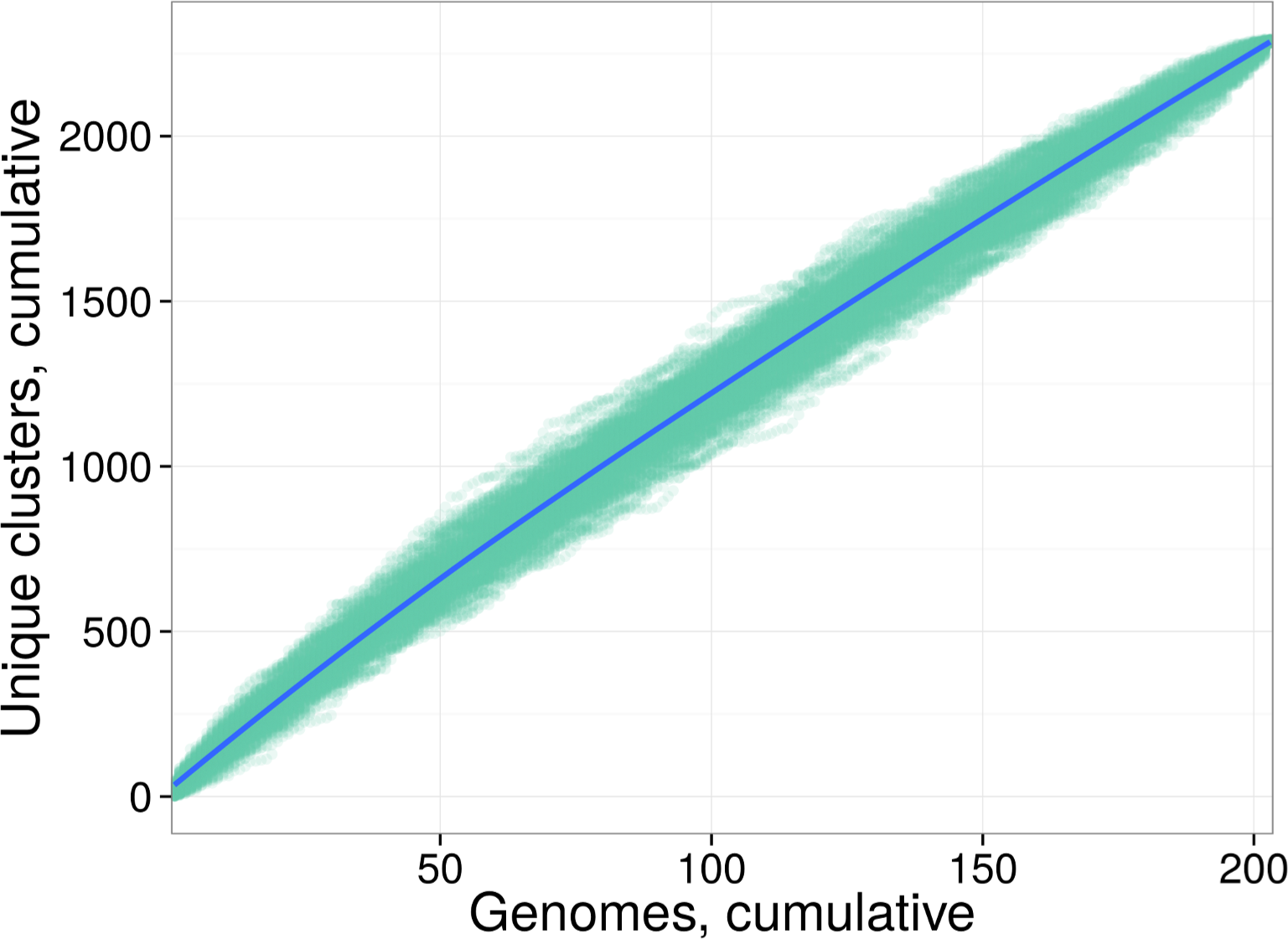
Total unique gene clusters smoothed scatterplot with genome order resampling. Similarity score threshold used: 0.5. Solid line is a loess fit to the scatterplot. The final point is always the same in all resamplings (the total number of unique gene clusters), causing scatterplot to narrow down at the last examined genome.

### Heap’s alpha

In studies of bacterial core and pan-genomes, the Heap’s alpha parameter is often used to estimate how many of the genes from a specific taxonomic group we haven’t seen yet. For pan-genomes, Heap’s alpha smaller than 1 means an open pan-genome (i.e. there are many more genes we haven’t seen yet), while alpha larger than 1 means a closed pan-genome (there is only a small/limited number of genes we haven’t seen yet). Rephrasing for our case, each genome has one or more groups of gene clusters (as described earlier), including duplicates. We applied the micropan R package [13] and obtained alpha = 0.129 for our set of genomes, indicating an open pan-clustome.

### Rarefaction analysis

Application of rarefaction analysis to extrapolate expected groups of gene clusters to higher genome counts has, in our opinion, argumentative value, because several assumptions of this analysis are likely not satisfied.

The first assumption is that “individuals are randomly distributed”. For BGC diversity, this implies that BGC types (and cluster similarity groups) are randomly distributed across genomes – which is not the case, as cluster composition is strongly defined by each bacteria’s environment, distinguishing, for example, clustomes of soil and water-dwelling bacteria. One has to control for the environment variable to satisfy this requirement.

The second assumption is that sample size is sufficiently large, which may or may not be the case.

Finally, to extrapolate from rarefaction analysis, one has to assume a certain underlying parent distribution.

On top of the technical limitations, rarefaction analysis does not account for several important biological peculiarities of BGCs. First of all, BGCs consist of biosynthetically-active sub-units (genes and gene modules), which can be re-arranged to produce different compounds, resulting in a combinatorial expansion of the BGCs diversity. Secondly, genome-based extrapolations rely solely on the information obtained from cultured strains. In other words, we might be missing a significant portion of BGCs diversity until methods are developed to grow hard to culture strains. Finally, genome-based analysis relies on the sequenced subset of bacterial genomes, inheriting biases which influenced strain selection for sequencing. As the number of genomes grows, this final bias should decrease.

That said, rarefaction analysis is still useful, and can be found in Appendix S4. The same analysis was performed on a small subset of 38 Streptomyces-only genomes; extrapolation indicated that Streptomyces alone might be responsible for approximately half of all the groups of BGCs we have yet to discover.

### Other types of analysis

There are several more types of analysis which can be performed on the table of similar gene cluster pairs generated by our software. Probably the first one to mention is looking for relationships between 16S rDNA phylogeny and BGC similarity, and/or percentage of unique clusters per genome. Succeeding in establishing a link between the two will provide more proof for the horizontal transfer of entire secondary metabolite BGCs. Another interesting (and related) question is whether fungal and bacterial BGCs have similar elements, and if yes – to what extent, and which types of BGCs? Our software should be applicable to fungal genomes as well, as long as BGC prediction will function for fungal genomes.

Another interesting type of analysis to perform is a graph of BGC similarity links (weighted edges) between all the genomes (nodes). This graph can be examined for links density, thus grouping genomes by BGC similarity. Another variation could add taxonomy information, e.g. by creating graph nodes from genus-level aggregates of genomes, with edges representing a sum of similarities between genomes of the connected genii. Described graph analysis may help identify biosynthetic hubs and evolutionary links between BGCs.

### AntiSMASH 3 known gene clusters report

AntiSMASH 3 (which has the functionality to report similarity to known gene clusters) was not yet available at the time of performing this study. As soon as it was published [14], we briefly compared our similarity approach to the known clusters reports of AntiSMASH 3.

The major difference is that AntiSMASH 3 is better suited for analysing individual genomes (or small groups thereof): BGCs of the given genome(s) are compared against a set of curated clusters. Conversely, our pipeline is better suited for the analysis of larger groups of genomes either with or without any reference BGC datasets, to produce overview BGC similarity statistics. Incremental analysis (comparing groups of new genomes to those analysed previously) is also possible.

There are also some other differences – for example, our protein similarity estimates are based on full-length alignments instead of protein BLAST results, and we use more cluster similarity metrics – but the most important difference is in the intended use.

### Limitations of the study

As secondary metabolite gene clusters were predicted using AntiSMASH 2, this study is clearly limited to the range of similarity-detectable compounds. However, assuming the same biosynthetic logic of possible new secondary metabolite BGCs predicted using other/newer methods, our approach to estimate BGC similarity and uniqueness should stay valid, and – thanks to the publicly available source code – can be applied to new prediction results as well.

Another potential pitfall was high internal similarity of the ∼300 genomes set downloaded from GenBank. However, we have invested significant effort into genome de-duplication (discarding ∼100 highly-similar genomes), and accounted for any remaining similar genome pairs during subsequent data analysis.

Finally, the number of currently available *draft* or otherwise *incomplete* Actinobacteria genomes is much higher than 300, but we intentionally limited the used genomes quality to *complete*, as otherwise, due to genome fragmentation, the number of predicted BGCs would be artificially inflated due to BGC fragment predictions.

### Future work

The approach (and software) that we present can be further improved with:

- better synteny measures, as discussed in *Gene-level synteny*;
- taking into account the order of biosynthetic domains (using a method similar to proper gene-level synteny); this should significantly increase precision of similarity measures;
- better gene cluster boundaries by the gene cluster prediction software; as discussed above, correct boundaries are crucial for proper similarity measurements;
- common genome annotation pipeline such as prokka [15] instead of stripping existing gene/CDS annotations and then performing basic re-annotation with Glimmer;
- possibly, predicted structure similarities as an additional metric; this improvement is hard to achieve, unless compound structure prediction from genes significantly improves;
- a graphical user interface.

We welcome interested groups to submit comments, suggestions, and patches.

### Applications

In addition to answering questions about gene clusters diversity, our software can be used to catalogue BGCs and create databases – either local, for one or several labs, or global, as a web-service. Implementing such a database, especially with incremental genome additions (which is already possible thanks to caching computationally-intensive steps) will allow further strengthening and improvement of genome-based BGC selection and activation approaches. An obvious use would be to focus on (extract and examine) only unique BGCs from a genome newly added to such a database. Another obvious use would be to look for similar BGCs (and additional producers) of the BGC under investigation. Even low similarity scores might be beneficial, as they help discovering BGCs with only some of the biosynthetic modules being similar, indicating a potential modified compound.

We have created and keep extending an internal database of similar BGCs for the purposes listed above. Further development should enable providing such a database service to the wider research community.

## Conclusions

We found that Actinobacteria (on average) have more predicted secondary metabolite gene clusters per genome than other bacteria. Cyanobacteria and Proteobacteria are runners-up in terms of the number of predicted gene clusters per genome.

The number of predicted gene clusters per genome in Actinobacteria correlates with genome size, increasing by almost 4 gene clusters per megabase beyond the first 1.7 megabases.

At the Genus level, Streptomyces, Frankia, Rhodococcus and Burkholderia have the highest average gene clusters per genome counts.

In the examined set of 3247 predicted secondary metabolite gene clusters from 208 de-duplicated Actinobacteria genomes we identified 40-48% as the upper boundary of the unique gene clusters percentage. Defining gene cluster similarity threshold (as well as compound [dis-]similarity) is a conceptually hard task – would a single gene make a difference, or two? What if this different gene results in a major compound modification? To formulate the similarity threshold, we used 4 pairs of known similar gene clusters and diagnostic plots across the full range of threshold values. We found that imprecise identification of gene cluster boundaries is the major hurdle for precise gene cluster similarity evaluation. On our dataset, Heap’s alpha is 0.129, and there is no saturation of the cumulative unique gene clusters curve for 208 genomes. This implies that Actinobacteria pan-clustome is open, and has many new gene clusters to be discovered.

Command-line software pipeline that we developed automates generation of the dataset which served as input for this article’s analysis. We also include scripts used for the analysis in the open git repository. Our software pipeline can be used to create and maintain databases of genomes and gene clusters, and to compare new genomes/clusters to such databases. As the next step, we plan developing such a database as an open service to the scientific community.

## Methods

### Secondary metabolite gene clusters

We use the term “secondary metabolite gene clusters” as it is used when publishing genome announcements. However, some of the secondary metabolite gene cluster (GC) types (siderophores being the most prominent example) are essential for bacteria survival. We decided to keep these BGC types in the study, so as not to artificially inflate the percentage of unique BGCs based on the currently-accepted use of “secondary”.

### Overview of bacterial biosynthetic potential

Genomes were downloaded on the 4th of February, 2014, from <u>ftp://ftp.ncbi.nlm.nih.gov/genomes/Bacteria/all.gbk.tar.gz</u>. The archive contained 2773 sub-directories, normally one directory per species. However, 9 directories contained multiple strain genomes of the same species, or even genomes of other species (S5 has a full list of such directories). For each of the genomes, multiple GenBank files (representing chromosomes and plasmids) were mechanically merged into a single multi-locus GenBank file. Table S6 lists all 5165 sequence accession numbers together with descriptions and sequence sizes. All 2775 genomes were then analysed with antiSMASH 2.0 [9]. For Actinobacteria-only biosynthetic potential overview, all 285 Actinobacteria results were copied out into a separate directory. For both the complete 2775-genome and 285-genome sets, results were collected into a CSV file using a custom summarize_antismash_results.py script (<u>https://bitbucket.org/qmentis/bioinformatics-scripts/src/b3faa96eeac2359a60016ded830ef3dab7c4f228/summarize_antismash_results.py</u>), and then loaded into R [16] for further analysis.

Both genome lists were searched for same species with different accessions. The highest number of such cases was observed for *Staphylococcus aureus subsp. aureus* ST228, from the genome evolution study [17]. After cleaning out highly similar genomes (see Appendix S7 for a table of accessions), 2724 genomes (278 for the Actinobacteria subset) were left for further analysis.

Genomes shorter than 2.5 megabases (Table S1) were very likely to have no or very few predicted BGCs; short genomes were removed from the all-bacteria overview, leaving 1651 genomes (Table S8) for analysis.

### Assessment of biosynthetic gene clusters diversity

285 complete Actinobacteria genomes from GenBank (see above) were supplemented with 25 genomes from other sources (all are Actinobacteria), then pruned from duplicates and highly-similar genomes (manual pruning, relying on the initial BGC comparison runs and on whole-genome Mauve [18] alignments), resulting in a list of 208 genomes. Secondary metabolite BGCs were predicted in these genomes using AntiSMASH 2 [9]. 5 of 208 genomes had zero predicted BGCs and were excluded from further analysis. Four pairs of known similar BGCs were then added to the analysis as controls and the basis for estimating BGC similarity thresholds (see Table S9 for a complete list of genomes used). See Fig. S10 for a flowchart of genomic data processing.

GC pair similarity is defined as a set of metrics, including count of similar gene pairs, average protein-level identity, synteny of cluster genes, and possibly others (Fig. 7). For simplicity, in this study overall BGC similarity score was defined as a ratio of genes similar between the clusters to the count of genes in those clusters, *score = 0.5 * (similar_genes/cluster1_genes + similar_genes/cluster2_genes*). Each BGC is categorized as “unique” or “similar to other clusters” based on the thresholds defined for each of the metrics. Minimal required protein-level identity was 60%; similar gene pairs were not reported below this identity. Synteny coefficient was calculated as gene order correlation coefficient: similar genes were sequentially numbered in both BGCs, followed by calculating Pearson correlation for both integer vectors. Downstream results analysis uncovered weak filtering capability of synteny coefficient, so despite being calculated it was not used for discerning similar and unique BGCs.

**Fig. 7.**
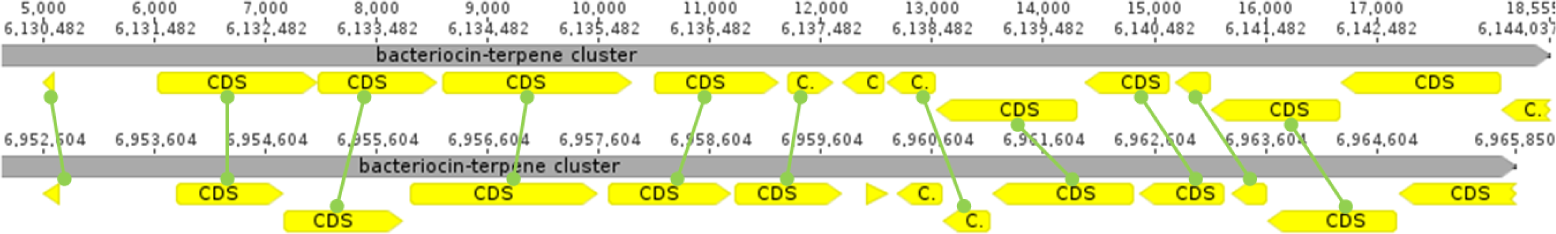
Example of a comparison of bacteriocin-terpene gene clusters from *Actinosynnema mirum* DSM 43827 (top) and *Saccharothrix espanaensis* DSM 44229 (bottom). Rounded-end handles between the CDS of the two genomes mark 11 gene pairs, where protein alignment identity was higher than 60%. For these 11 gene pairs, synteny coefficient (gene order correlation) is 1.0, and average actual protein identity is 82%. Gene cluster similarity score is 0.76.

Four pairs of known BGCs from different genomes responsible for the production of the same compound (landomycin, microsclerodermin, moenomycin, nonactin) were used as controls.

### Cumulative unique gene clusters curve with genome order resampling

To look for possible saturation of the total number of unique gene clusters from examined genomes, we performed the following steps:

1. Randomize genome order.
2. Take the 1^st^ genome. As it is the first seen genome, all of its BGCs are unique. Plot the number of observed BGCs (X: genome number; Y: total unique BGCs found).
3. Take the 2^nd^ genome, compare its BGCs to those seen before. Sum unseen (unique) BGCs from the 2^nd^ genome with the unique BGCs from the 1^st^, and plot the cumulative value.
4. Repeat until all the genomes are examined, every time comparing genome’s clusters only to those we had seen in previously examined genomes. At every genome, plot cumulative unique BGCs.
5. Reset seen BGCs to an empty set. Repeat steps 1-4 200 times.

### Grouping gene clusters by similarity

Sample gene clusters A, B, and C will be put into a single group if they have pairwise links with similarity scores above the specified threshold. For example, high-scoring links A-B and B-C are sufficient to put all three into a single group, even if A-C happens to be below the threshold. Alternatively, A-B and A-C are sufficient to group all three, even if B-C is below the threshold. Unique BGCs become singleton groups.

### Rarefaction analysis

Rarefaction analysis was performed in both incidence and abundance modes using the iNEXT R package [19]. Abundance analysis was performed on counts of gene clusters (“individuals”) per group of gene clusters (“species”, see *Grouping gene clusters by similarity* above), extrapolating to 45 and 60 thousand examined gene clusters in total. Incidence analysis was performed on a vector of genome counts (“samples”) per group of gene clusters (“species”), extrapolating to 15 thousand genomes.

## Supporting information captions

**S1 Table. Summary of 1074 deduplicated small genomes shorter than 2.5 megabases**.

**S2 Appendix. Diagnostic plots for ranges of threshold values**.

**S3 Appendix. Total unique gene clusters resampling scatterplots**.

**S4 Appendix. Rarefaction plots**.

**S5 Text. Multigenome directories downloaded from GenBank FTP**.

**S6 Table. Table of 2775 genomes used for overview analysis**.

**S7 Appendix. List of highly similar genomes removed prior to analysis**.

**S8 Table. 1651 deduplicated genomes larger than 2.5 megabases used for overview analysis**.

**S9 table. All 318 genomes and sequences used for similarity analysis**.

**S10 Figure. Similarity analysis genomic data processing flowchart**.

**S11 Dataset. Full table (tab-separated values) of similar gene cluster pairs**.

